# Recurrent convolutional neural networks: a better model of biological object recognition

**DOI:** 10.1101/133330

**Authors:** Courtney J. Spoerer, Patrick McClure, Nikolaus Kriegeskorte

**Affiliations:** Medical Research Council Cognition and Brain Sciences Unit, University of Cambridge, 15 Chaucer Road, Cambridge, CB2 7EF, United Kingdom

## Abstract

Feedforward neural networks provide the dominant model of how the brain performs visual object recognition. However, these networks lack the lateral and feedback connections, and the resulting recurrent neuronal dynamics, of the ventral visual pathway in the human and nonhuman primate brain. Here we investigate recurrent convolutional neural networks with bottom-up (B), lateral (L), and top-down (T) connections. Combining these types of connections yields four architectures (B, BT, BL, and BLT), which we systematically test and compare. We hypothesized that recurrent dynamics might improve recognition performance in the challenging scenario of partial occlusion. We introduce two novel occluded object recognition tasks to test the efficacy of the models, *digit clutter* (where multiple target digits occlude one another) and *digit debris* (where target digits are occluded by digit fragments). We find that recurrent neural networks outperform feedforward control models (approximately matched in parametric complexity) at recognising objects, both in the absence of occlusion and in all occlusion conditions. Recurrent networks were also found to be more robust to the inclusion of additive Gaussian noise. Recurrent neural networks are better in two respects: (1) they are more neurobiologically realistic than their feedforward counterparts; (2) they are better in terms of their ability to recognise objects, especially under challenging conditions. This work shows that computer vision can benefit from using recurrent convolutional architectures and suggests that the ubiquitous recurrent connections in biological brains are essential for task performance.

## 1 Introduction

The primate visual system is highly efficient at object recognition, requiring only brief presentations of the stimulus to perform the task (Potter, 1976; Thorpe et al., 1996; Keysers et al., 2001).

Within 150ms of stimulus onset, neurons in inferior temporal cortex (IT) encode object information in a form that is robust to transformations in scale and position (Hung et al., 2005; Isik et al., 2014), and is predictive of human behavioural responses (Majaj et al., 2015).

This rapid processing lends support to the idea that invariant object recognition can be explained through a feedforward process (DiCarlo et al., 2012), a claim that has been supported by the recent successes of feedforward neural networks in computer vision (e.g. Krizhevsky et al., 2012) and the usefulness of these networks as models of primate visual processing (Riesenhuber and Poggio, 1999; Serre et al., 2007; Wallis and Rolls, 1997; Yamins et al., 2013, 2014; Khaligh-Razavi and Kriegeskorte, 2014; Güçlü and van Gerven, 2015).

The success of feedforward models of visual object recognition has resulted in feedback processing being underexplored in this domain. However, both anatomical and functional evidence seems to suggest that feedback connections play a role in object recognition. For instance, it is well known that the ventral visual pathway contains similar densities of feedforward and feedback connections (Felleman and Van Essen, 1991; Sporns and Zwi, 2004; Markov et al., 2014), and functional evidence from primate and human electrophysiology experiments show that processing of object information unfolds over time, beyond what would be interpreted as feedforward processing (Sugase et al., 1999; Brincat and Connor, 2006; Freiwald and Tsao, 2010; Carlson et al., 2013; Cichy et al., 2014; Clarke et al., 2015). Some reports of robust object representations, normally attributed to feedforward processing (Isik et al., 2014; Majaj et al., 2015), occur within temporal delays that are consistent with fast local recurrent processing (Wyatte et al., 2014). This suggests that we need to move beyond the standard feedforward model if we are to gain a complete understanding of visual object recognition within the brain.

Fast local recurrent processing is temporally dissociable from attentional effects in frontal and parietal areas, and is thought to be particularly important in recognition of the degraded objects (for a review see Wyatte et al., 2014). In particular, object recognition in the presence of occlusion is thought to engage recurrent processing. This is supported by the finding that recognition under these conditions produces delayed behavioural and neural responses, and recognition can be disrupted by masking, which is thought to interfere with recurrent processing (Johnson and Olshausen, 2005; Tang et al., 2014; Wyatte et al., 2012). Furthermore, competitive processing, which is thought to be supported by lateral recurrent connectivity (Adesnik and Scanziani, 2010), aids recognition of occluded objects (Kolankeh et al., 2015). Scene information can also be decoded from areas of early visual cortex that correspond to occluded regions of the visual field (Smith and Muckli, 2010) further supporting the claim that feedback processing is engaged when there is occlusion in the visual input.

Occluded object recognition has been investigated using neural network models in previous work, which found an important role for feedback connections when stimuli were partially occluded (O'Reilly et al., 2013). However, the type of occlusion used in these simulations, and previous experimental work, has involved fading out or deleting parts of images (Wyatte et al., 2012; Smith and Muckli, 2010; Tang et al., 2014). This does not correspond well to vision in natural environments where occlusion is generated by objects occluding one another. Moreover, deleting parts of objects, as opposed to occluding them, leads to poorer accuracies and differences in early event-related potentials (ERPs) that could indicate different effects on local recurrent processing (Johnson and Olshausen, 2005). Therefore, it is important to investigate the effects of actual object occlusion in neural networks to complement prior work on deletion.

In scenes where objects occlude one another it is important to correctly assign border ownership for successful recognition. Border ownership can be thought of as indicating which object is the occluder and which object is being occluded. Border ownership cells require information from outside their classical receptive field and border ownership signals are delayed relative to the initial feedforward input, which both suggest the involvement of recurrent processing (Craft et al., 2007). A number of computational models have been developed to explain border ownership cells. What is common amongst these models is the presence of lateral or top-down connections (Zhaoping, 2005; Sakai and Nishimura, 2006; Craft et al., 2007). The importance of recurrent processing for developing selectivity to border ownership further suggests that recurrence has an important role for recognising occluded objects.

To test the effects of occlusion, we developed a new generative model for occlusion stimuli. The images contain parameterised, computer-generated digits in randomly jittered positions (optionally, the size and orientation can also be randomly varied). The code for generating these images is made available at https://github.com/cjspoerer/digitclutter. The task is to correctly identify these digits. Different forms of occlusion are added to these images, including occlusion from non-targets and other targets present in the image, we refer to these as *digit debris* and *digit clutter*, respectively. The first form of occlusion, digit debris, simulates situations where targets are occluded by other objects that are task irrelevant. The second case, digit clutter, simulates occlusion where the objective is to account for the occlusion without suppressing the occluder, which is itself a target. This stimulus set has a number of benefits. Firstly, the underlying task is relatively simple to solve, which allows us to study the effects of occlusion and recurrence with small-scale neural networks. Therefore, any challenges to the network will only result from the introduction of occlusion. Additionally, as the stimuli are procedurally generated, they can be produced in large quantities, which enables the training of the networks.

Recurrent processing is sometimes thought of as cleaning up noise, where occlusion is a special case of noise. A simple case of noise is additive Gaussian noise, but we hypothesise that recurrence is unlikely to show benefits in these conditions. Consider the case of detecting simple visual features that show no variation, e.g. edges of different orientations. An optimal linear filter can be learnt for detecting these features. This linear filter would remain optimal under independent, additive Gaussian noise, as the expected value of the input and output will remain the same under repeated presentations. Whilst this result does not exactly hold for the case of non-linear filters that are normally used in neural networks, we might expect similar results. Therefore, we would expect no specific benefit of recurrence in the presence of additive Gaussian noise. If this is true, we can infer that the role of recurrence is not for performing object recognition in noisy conditions, generally. Otherwise, it would support the conclusion that reccurence is useful across a wider range of challenging conditions.

In this work, we investigate object recognition using convolutional neural networks. We extend the idea of the convolutional architecture to networks with bottom-up (B), lateral (L), and top-down (T) connections in a similar fashion to previous work (Liang and Hu, 2015; Liao and Poggio, 2016). These connections roughly correspond to processing information from lower and higher regions in the ventral visual hierarchy (bottom-up and top-down connections), and processing information from within a region (lateral connections). We choose to use the convolutional architecture as it is a parameter efficient method for building large neural networks that can perform real-world tasks (LeCun et al., 2015). It is directly inspired by biology, with restricted receptive fields and feature detectors that replicate across the visual field (Hubel and Wiesel, 1968) and advances based on this architecture have produced useful models for visual neuroscience (Kriegeskorte, 2015). The interchange between biology and engineering is important for the progress of both fields (Hassabis et al., 2017). By using convolutional neural networks as the basis of our models, we aim to maximise the transfer of knowledge from these more biologically motivated experiments to applications in computer vision, and by using recurrent connections, we hope that our models will contribute to a better understanding of recurrent connections in biological vision whilst maintaining the benefits of scalability from convolutional architectures.

To test whether recurrent neural networks perform better than feedforward networks at occluded object recognition, we trained and tested a range of networks to perform a digit recognition task under varying levels of occlusion. Any difference in performance reflects the degree to which networks learn the underlying task of recognising the target digits, and handle the occlusion when recognising the digit. To differentiate between these two cases we also look at how well networks trained on occluded object recognition generalise to object recognition without occlusion. We also test whether recurrence shows an advantage for standard object recognition and when dealing with noisy inputs, more generally, by measuring object recognition performance with and without the presence of additive Gaussian noise. Finally, we study whether any benefit of recurrence extends to occluded object recognition where the occluder is also a target, the networks are tested on multiple digit recognition tasks where the targets overlap.

## 2 Material and Methods

### 2.1 Generative model for stimuli

To investigate the effect of occlusion in object recognition, we opt to use a task that could be solved trivially without the presence of occlusion, computer generated digit recognition. Each digit uses the same font, colour, and size. The only variable is the position of the digit, which is drawn from a uniform random distribution. This means, the only invariance problem that needs to be solved is translation invariance, which is effectively built into the convolutional networks we use. Therefore, we restrict ourselves to only altering the level of occlusion to increase task difficulty. This means we need to use some challenging occlusion scenarios to differentiate between the models. However, this allows us to isolate the effects of occlusion and, by keeping the overall task relatively simple, we can use small networks, allowing us to train them across a wide range of conditions.

We generate occlusion using two methods, by scattering debris across the image, digit debris, and by presenting overlapping digits within a scene, which the network has to simultaneously recognise, digit clutter.

For digit debris, we obtain debris from fragments of each of the possible targets, taking random crops from randomly selected digits. Each of these fragments are then added to a mask that is overlaid on the target digit (Figure 1). As a result, the visual features of non-target objects, that the network has to ignore, are present in the scene. These conditions mean that summing the overall visual features present for each digit becomes a less reliable strategy for inferring the target digit. This is in contrast to deletion where there is only a removal of features that belong to the target.

**Figure 1:**
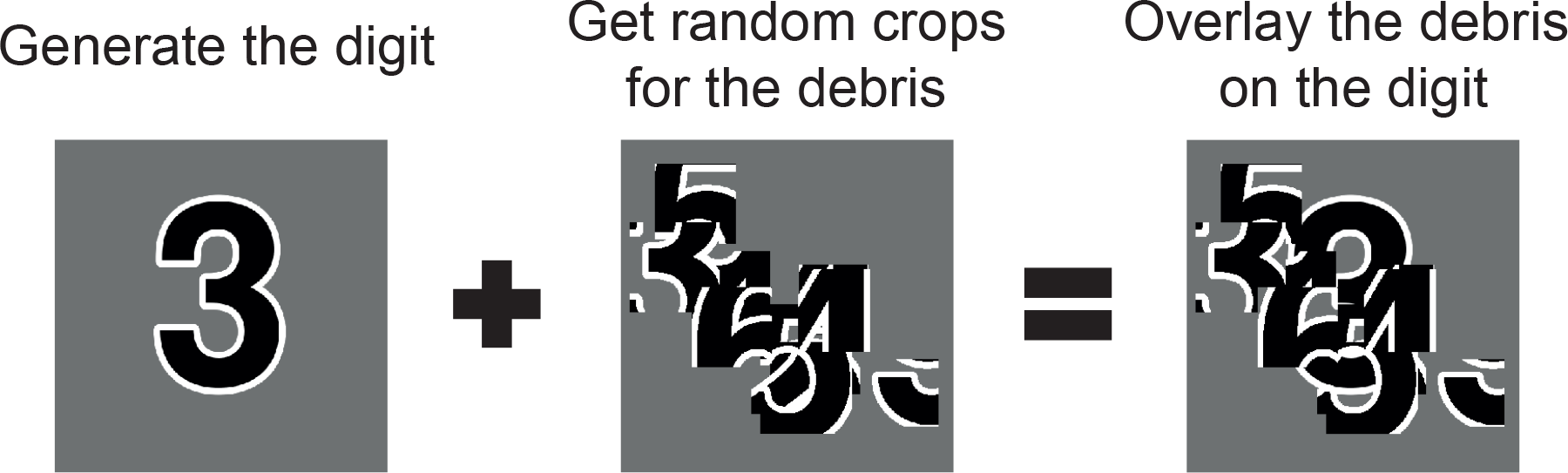
The process for generating stimuli for digit debris. First the target digit is generated. Random crops of all possible targets are taken to create a mask of debris, which is applied to the target as an occluder.

However, within natural visual scenes, occlusion is generated by other whole objects. These objects might also be of interest to the observer. In this scenario, simply ignoring the occluding objects would not make sense. In digit clutter, these cases are simulated by generating images with multiple digits that are sequentially placed in an image, where their positions are also drawn from a uniform random distribution. This generates a series of digits that overlap, producing a relative depth order. The task of the network is then to recognise all digits that are present.

Design of these images was performed at high resolution (512 × 512 pixels) and, for computational simplicity, the images were resized to a low resolution (32 × 32 pixels) when presented to the network.

In these experiments we use stimulus sets, that vary in either the number of digits in a scene - three digits, four digits, or five digits - or the number of fragments that make up the debris - 10 fragments (light debris), 30 fragments (moderate debris), or 50 fragments (heavy debris). Examples from these stimulus sets are shown in Figure 2. This allows us to measure how the performance of the networks differ across these task types and levels of occlusion.

**Figure 2:**
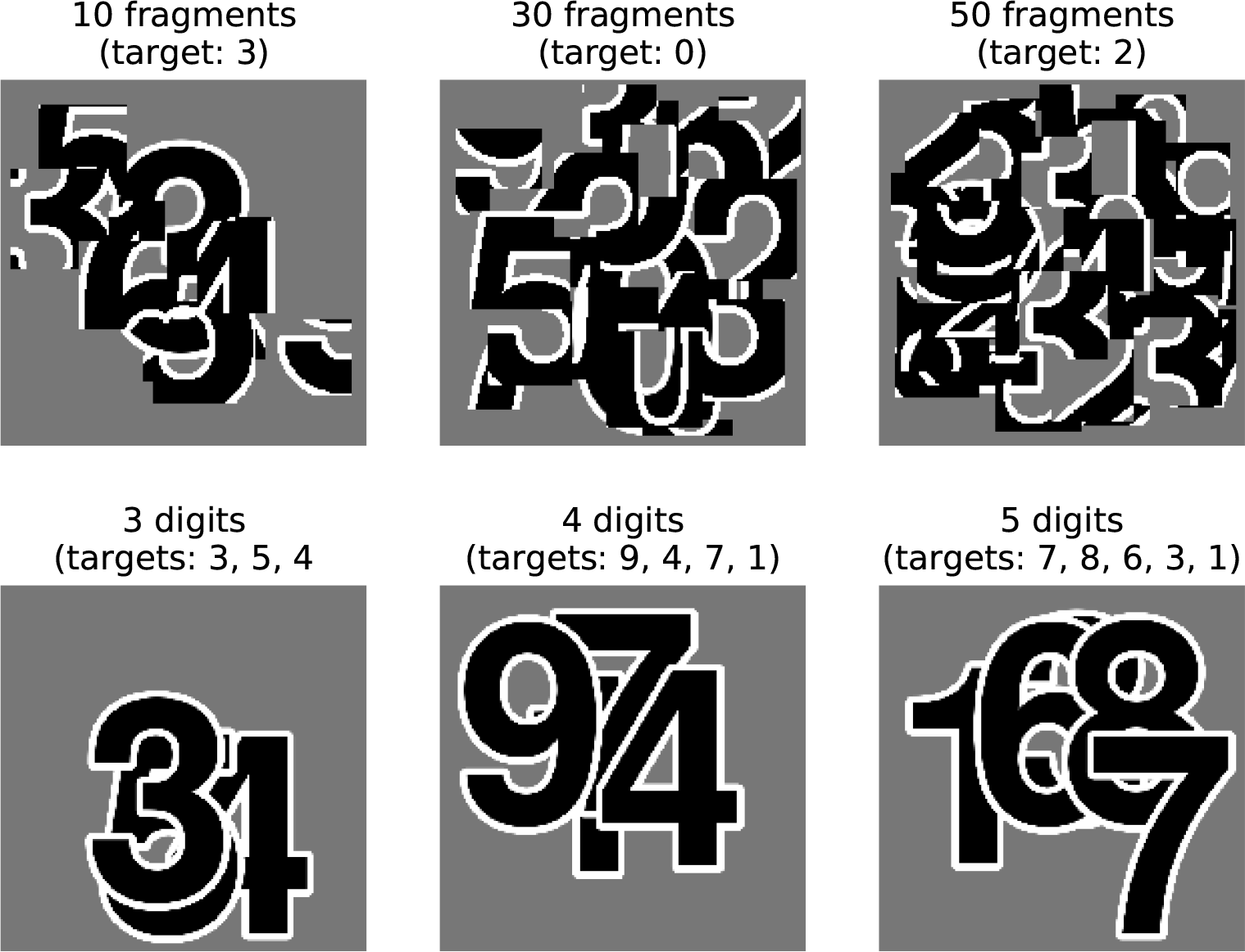
High resolution examples from the stimulus sets used in these experiments. The top row shows digit debris stimuli for each of the three conditions tested here, with 10, 30, and 50 fragments. The bottom row shows digit clutter stimuli with 3, 4, and 5 digits.

For each of these image sets, we randomly generated a training set of 100,000 images and a validation set of 10,000 images, which were used for the determining the hyperparameters and learning regime. All analyses where performed on an independent test set of 10,000 images.

All images underwent pixel-wise normalisation prior to being passed to the network. For an input pixel *x* in position *i, j*, this is defined as

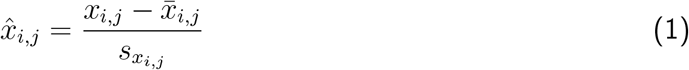

where *x_i,j_* is the raw pixel value, *x*̅*_i,j_* is the mean pixel value and *s_x_i,j__* is the standard deviation of pixel values. The mean and standard deviation are computed for each specific position across the whole of the training data.

To test the hypothesis that the benefit of recurrence is not simply for cleaning up noise, we also test the network on object recognition where the input has additive Gaussian noise. To prevent ceiling performance, we use the MNIST handwritten digit recognition data set (LeCun et al., 1998). The MNIST data set contains 60,000 images in total that are divided into a training set of 50,000 images, a validation set of 5,000 images, and a testing set of 10,000 images.

We add Gaussian noise to these images after normalisation, which allows an easy interpretation in terms of signal to noise ratio. In this case, we use Gaussian noise with a standard deviation of 1 and 2, which produces images with a signal-to-noise ratio (SNR) of 1 and 0.5, respectively.

### 2.2 Models

In these experiments we use a range of convolutional neural networks (for an introduction to this architecture, see Goodfellow et al., 2016). These networks can be categorised by the particular combination of bottom-up, lateral, and top-down connections that are present. As it does not make sense to construct the networks without bottom-up connections (as information from the input cannot reach higher layers), we are left with four possible architectures with the following connections, bottom-up only (B), bottom-up and top-down (BT), bottom-up and lateral (BL), and bottom-up, lateral and top-down (BLT). Each of these architectures are illustrated schematically in Figure 3.

**Figure 3:**
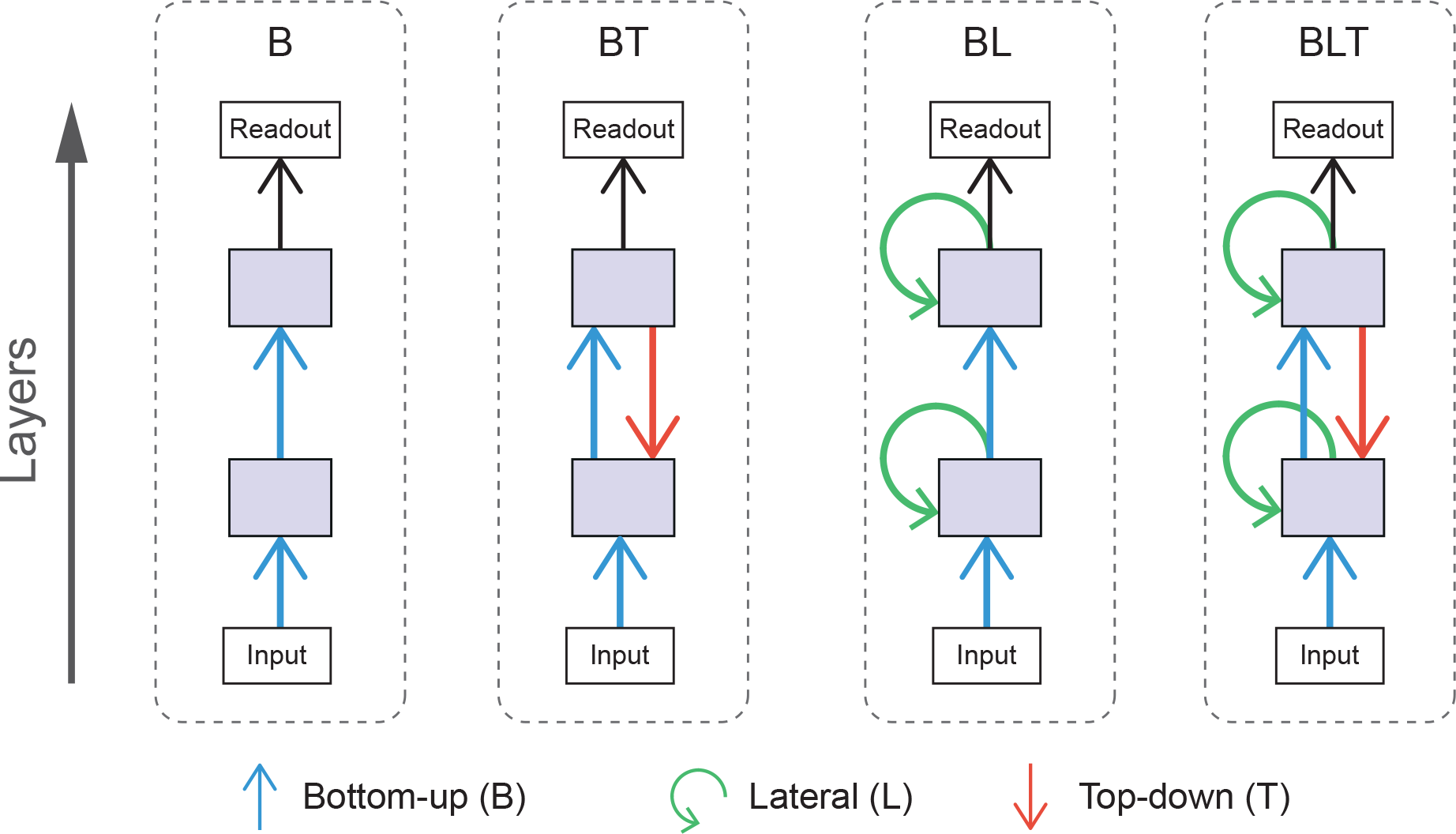
Schematic diagrams for each of the architectures used. Arrows indicate bottom-up (blue), lateral (green), and top-down (red) convolutions.

Adding top-down or lateral connections to feedforward models introduces cycles into the graphical structure of the network. The presence of cycles in these networks allow recurrent computations to take place, introducing internally generated temporal dynamics to the models. In comparison, temporal dynamics of feedforward networks can only be driven by changes in the input. The effect of recurrent connections can be seen through the unrolling of the computational graph across time steps. In these experiments, we run our models for four time steps and the resulting graph for BLT is illustrated in Figure 4.

**Figure 4:**
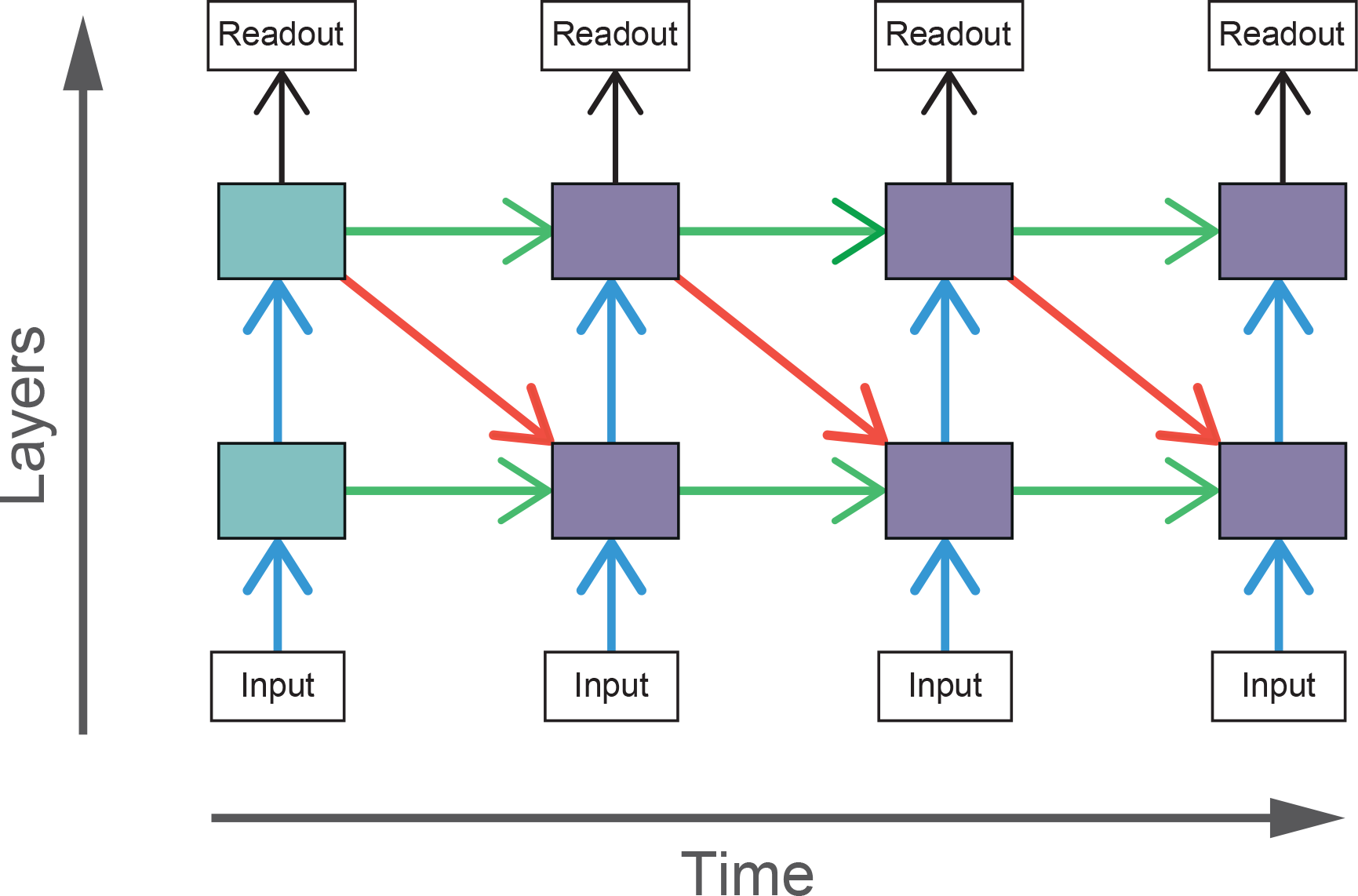
The computational graph of BLT unrolled over four time steps. The shaded boxes indicate hidden layers that receive purely feedforward input (blue) and those that receive both feedforward and recurrent input (purple).

As the recurrent networks (BT, BL, and BLT) have additional connections compared to purely feedforward networks (B), they also have a larger number of free parameters (Table 1). To control for this difference, we test two variants of B that have a more similar number parameters to the recurrent networks. The first control increases the number of features that can be learned by the bottom-up connections and the second control increases the size of individual features (known as the kernel size). These are referred to as B-F and B-K, respectively. Conceptually, B-K is a more appropriate control compared to B-F, as it effectively increases the number of connections that each unit has, holding everything else constant. In comparison, B-F increases the number of units within a layer, altering the layers representational power, in addition to changing the number of parameters. However, B-F is more closely parameter matched to some of the recurrent models, which motivates the inclusion of B-F in our experiments.

**Table 1:** Brief descriptions of the models used in these experiments including the number of learnable parameters and the number of units in each model.

| Model | Kernel size | No. Features | No. parameters | No. units |
| --- | --- | --- | --- | --- |
| B | $3 \times 3$ | 32 | 9,920 | 40,970 |
| B-F | $3 \times 3$ | 64 | 38,272 | 81,930 |
| B-K | $5 \times 5$ | 32 | 26,816 | 40,970 |
| BT | $3 \times 3$ | 32 | 19,168 | 40,970 |
| BL | $3 \times 3$ | 32 | 28,416 | 40,970 |
| BLT | $3 \times 3$ | 32 | 37,568 | 40,970 |

### 2.2.1 Architecture overview

All of the models tested consist of two hidden recurrent convolutional layers (described in Section 2.2.2) followed by a readout layer (described in Section 2.2.3). Bottom-up and lateral connections are implemented as standard convolutional layers with a 1×1 stride. The feedforward inputs between the hidden layers go through a max pooling operation, with a 2×2 stride and a 2×2 kernel. This has the effect of reducing the height and width of a layer by a factor of two. As a result, we cannot use standard convolutions for top-down connections, as the size of the top-down input from the second hidden layer would not match the size of the first hidden layer. To increase the size of the top-down input, we use transposed convolution (also known as deconvolution Zeiler et al., 2010) with an output stride of 2×2. This deconvolution increases the size of the top-down input so that it matches the size of the first hidden layer. The connectivity of this layer can be understood as a normal convolutional layer with 2×2 stride where the input and output sides of the layer have been switched.

As feedforward networks do not have any internal dynamics and the stimuli are static, feedforward networks only run for one time step. Each of the recurrent networks are run for four time steps. This is implemented as a computational graph unrolled over time (Figure 4), where the weights for particular connections are shared across each time step. The input is also replicated at each time point.

To train the network, error is backpropagated through time for each time point (Section 2.2.4), which means that the network is trained to converge as soon as possible, rather than at the final time step. However, when measuring the accuracy, we use the predictions at the final time step as this generally produces the highest accuracy.

### 2.2.2 Recurrent convolutional layers

The key component of these models is the recurrent convolutional layer (RCL). The inputs to these layers are denoted by ***h***_(*τ,m,i,j*)_, which represents the vectorised input from a patch centred on location *i, j*, in layer *m*, computed at time step *τ*, across all features maps (indexed by *k*). We define ***h***_(*τ*,0,*i,j*)_ as the input image to the network.

For B, the lack of recurrent connections reduces RCLs to a standard convolutional layer where the pre-activation at time step *τ* for a unit in layer *m*, in feature map *k*, in position *i, j* is defined as

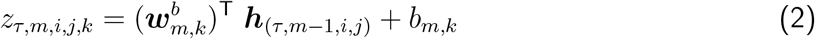

where *τ* = 0 (as B only runs for a single time step) the convolutional kernel for bottom-up connections is given in vectorised format by 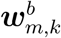 and the bias for feature map *k* in layer *m* is given by *bm,k*.

In BL, lateral inputs are added to the pre-activation, giving

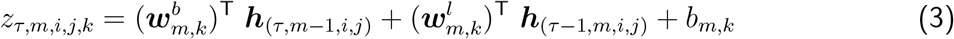

The term for lateral inputs 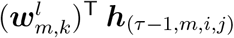 uses the same indexing conventions as the bottom-up inputs in Equation 2, where 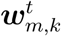 is the lateral convolutional kernel in vectorised format. As the lateral input is dependent on outputs computed on the timestep *τ* − 1, they are undefined for the first time step, when *τ* = 0. Therefore, when *τ* = 0 we set recurrent inputs to be a vector of zeros. This rule applies for all recurrent input, including top-down inputs.

In BT, we add top-down inputs to the pre-activation instead of lateral inputs. This gives

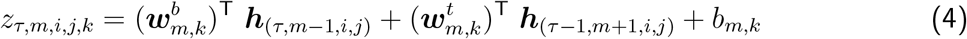

Where the top-down term is 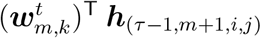, and 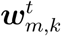 is the top-down convolutional kernel in vectorised format. In our models, top-down connections can only be received from other hidden layers. As a result, top-down inputs are only given when *m* = 1 and otherwise they are set to a vector of zeros. The rule for top-down inputs also applies to top-down inputs in BLT.

Finally, we can add both lateral and top-down inputs to the pre-activation, which generates the layers we use in BLT

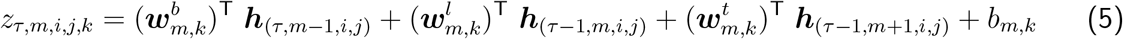

The output, *h_τ,m,i,j,k_*, is calculated using the same operations for all layers. The pre-activation *z_τ,m,i,j,k_* is passed through a layer of rectified linear units (ReLUs), and local response normalization (Krizhevsky et al., 2012).

ReLUs are defined as

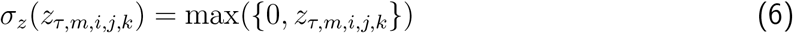

and local response normalisation is defined for input *x_τ,m,i,j,k_* as

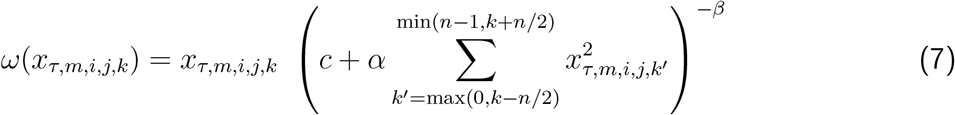

For local response normalisation, we use *n* = 5, *c* = 1, *α* = 10^*−*4^, and *β* = 0.5 throughout. This has the effect of inducing competition across the *n* closest features within a spatial location. The features are ordered arbitrarily and this ordering is held constant.

The output of layer *l* at time step *t* is then given by

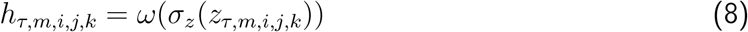

### 2.2.3 Readout layer

In the final layer of each time step, a readout is calculated for each class. This is performed in three steps. The first stage is a global max pooling layer, which returns the maximum output value for each feature map. The output of the global max pooling layer is then used as input to a fully connected layer with 10 output units. These outputs are passed through a sigmoid non-linearity, *σ_y_* (*x*), defined as

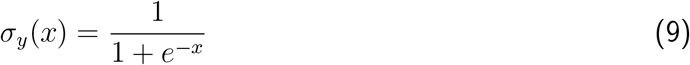

This has the effect of bounding the output between 0 and 1. The response of each of these outputs can be interpreted as the probability that each target is present or not.

### 2.2.4 Learning

At each time step, the networks give an output from the readout layer, which we denote ***y***̂*_t_*, where we interpret each output as the probability that a particular target is present or not.

In training, the objective is to match this output to a ground truth ***y***, which uses binary encoding such that its elements *y_i_* are defined as

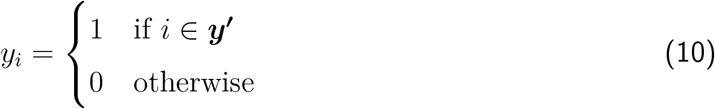

Where ***y′*** is the list of target digits present.

We used cross-entropy to calculate the error between ***y***̂*_t_* and ***y***, which is summed across all time steps

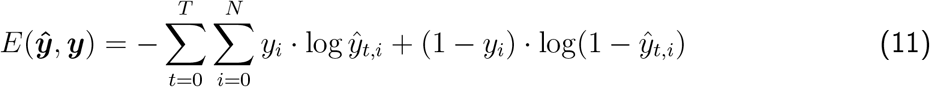

L2-regularisation is included, with a coefficient of *λ* = 0.0005, making the overall loss function

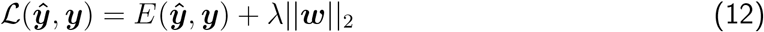

Where ***w*** the vector of all trainable parameters in the model.

This loss function was then used to train the networks by changing the parameters at the end of each mini-batch of 100 images according to the momentum update rule

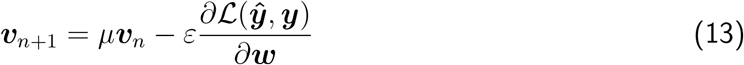

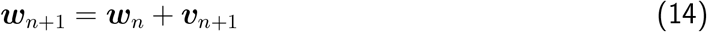

Where *n* is the iteration index, *µ* is the momentum coefficient, and *ε* is the learning rate. We use *µ* = 0.9 for all models and set *ε* by the following weight decay rule

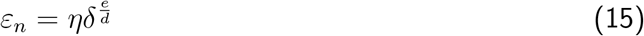

Where *η* is the initial learning rate, *δ* is the decay rate, *e* is the epoch (a whole iteration through all training images), and *d* is the decay step. In our experiments we use *η* = 0.1, *δ* = 0.1, and *d* = 40. All networks were trained for 100 epochs. The parameters for the training regime where optimized manually using the validation set.

## 2.3 Analysing model performance

### 2.3.1 Comparing model accuracy

We measured the performance of the networks by calculating the accuracy across the test set. For digit clutter tasks with multiple labels, we took the top-*n* class outputs as the network predictions, where *n* is the number of digits present in that task.

Accuracy was compared within image sets by performing pairwise McNemar's tests between all of the trained models (McNemar, 1947). McNemar's test is used here, which uses the variability in performance across stimuli as the basis for statistical inference (Dietterich, 1998). This does not require repeated training from different random seeds, which is both computationally expensive, and redundant, as networks converge on highly similar performance levels. By avoiding the need to retrain networks from different random initialisations we are able to explore a variety of qualitatively different architectures and infer differences between them.

To mitigate the increased risk of false positives due to multiple comparisons, we control the false discovery rate (the expected proportion of false positives among the positive outcomes) at 0.05 for each group of pairwise tests using the Benjamini-Hochberg procedure (Benjamini and Hochberg, 1995).

### 2.3.2 Comparing model robustness

To understand whether networks have varying levels of robustness to increased task difficulty (i.e. increased levels of debris, clutter, and Gaussian noise), we test for differences in the increase in error between all networks as task difficulty increases.

To achieve this, we fit a linear model to the error rates for each network separately, with the difficulty levels as predictors (e.g. light debris = 1, moderate debris = 2, heavy debris = 3). We extract the slope parameters from the linear models for a pair of networks and test if the difference in these slope parameters significantly differs from zero, by using a permutation test.

To construct a null distribution for the permutation test, we randomly shuffle predictions for a single image between a pair of networks. Error rates are then calculated for these shuffled predictions. A linear model is fit to these sampled error rates, for each model separately, and the difference between the slope parameters is entered into the null distribution. This procedure is run 10,000 times to approximate the null distribution. The *p*-value for this test is obtained by making a two-tailed comparison between the observed value for the difference in slope parameters and the null distribution. Based on the uncorrected *p*-values, a threshold is chosen to control the FDR at 0.05.

## 3 Results

### 3.1 Recognition of digits under debris

#### 3.1.1 Learning to recognise digits occluded by debris

Networks were trained and tested to recognise digits under debris to test for a particular benefit of recurrence when recognising objects under structured occlusion. We used three image sets containing different levels of debris, 10 fragments (light debris), 30 fragments (moderate debris), and 50 fragments (heavy debris). For every model, the error rate was found to increase as the level of debris in the image increased (Figure 5).

Under light and moderate debris, all but one of the pairwise differences were found to be significant (FDR = 0.05) with no significant difference between BL or BLT for light (*χ*2(1, N = 10, 000) = 0.04, *p* = 0.835) and moderate debris (*χ*^2^(1, *N* = 10, 000) = 0.00, *p* = 0.960). Of the feedforward models, B-K was the best performing. The error rates for each of the models are shown in Table 2.

**Figure 5:**
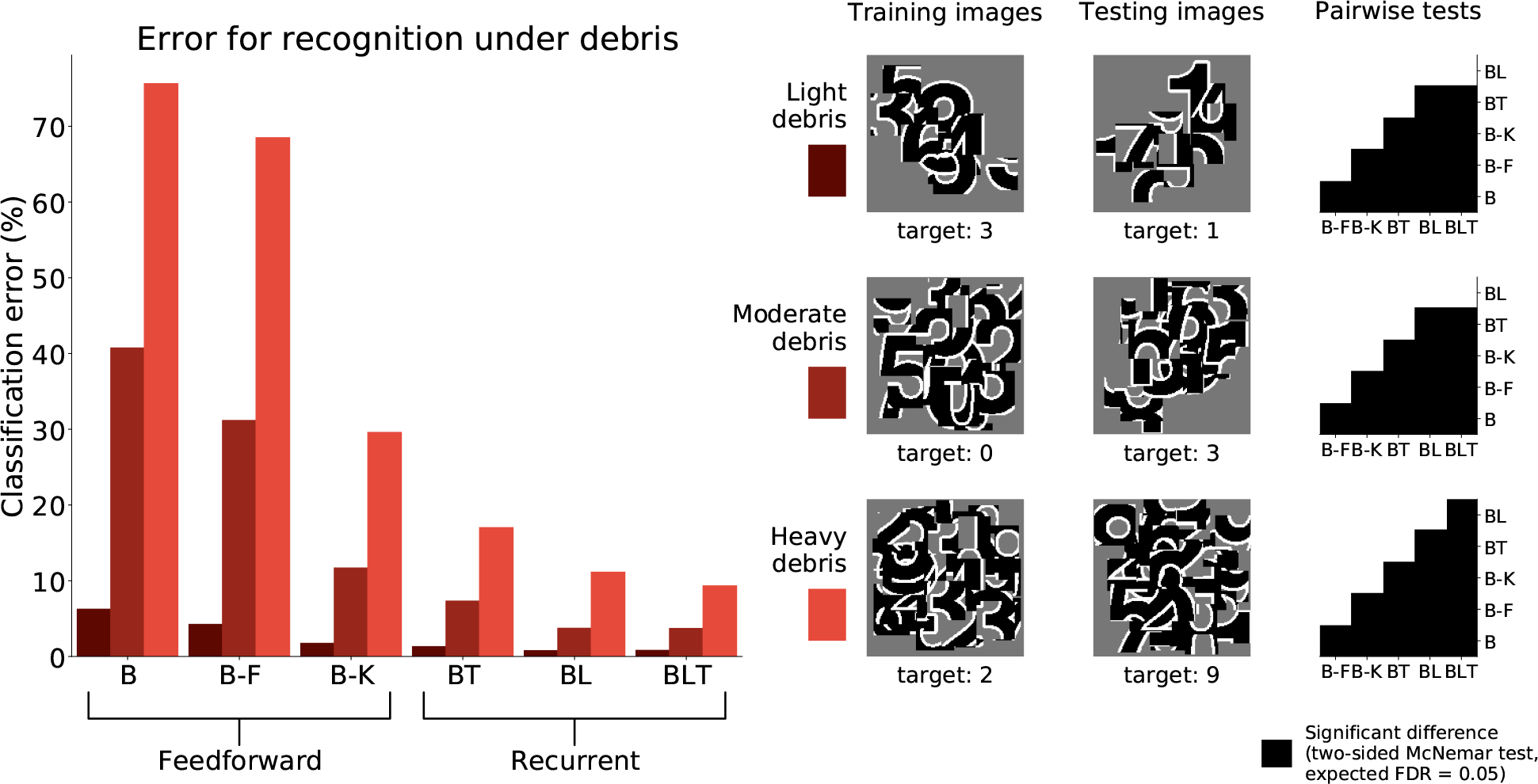
Classification error for all models on single digit detection under varying levels of debris. Examples of the images used to train and test the networks are also shown. Matrices to the right indicate significant results of pairwise McNemar tests. Comparisons are across models and within image sets. Black boxes indicate significant differences at p < 0.05 when controlling the expected false discovery rate at 0.05.

**Table 2:** Classification error for all of the models on single digit detection with varying levels of debris.

| Image set | B | B-F | B-K | BT | BL | BLT |
| --- | --- | --- | --- | --- | --- | --- |
| Light debris | 6.24% | 4.23% | 1.73% | 1.30% | 0.77% | 0.80% |
| Moderate debris | 40.73% | 31.16% | 11.68% | 7.31% | 3.72% | 3.70% |
| Heavy debris | 75.63% | 68.49% | 29.58% | 17.01% | 11.13% | 9.32% |

Under heavy debris all pairwise differences were significant (FDR = 0.05) including the difference between BL and BLT, which was not significant under light and moderate debris, with BLT out-performing BL. This suggests that at lower levels of occlusion, feedforward and lateral connections are sufficient for good performance. However, top-down connections become beneficial when the task involves recognising digits under heavier levels of debris.

#### 3.1.2 Learning to recognise unoccluded digits when trained with occlusion

To test if the networks learn a good model of the digit when trained to recognise the digit under debris, we test the performance of networks when recognising unoccluded digits.

When networks were trained to recognise digits under heavy debris, and tested to recognise unoccluded digits, we found all pairwise differences to be significant (FDR = 0.05, Figure 6). The best performing network was B-K, followed by recurrent networks. B and B-F performed much worse than all of the other networks (Table 3).

**Figure 6:**
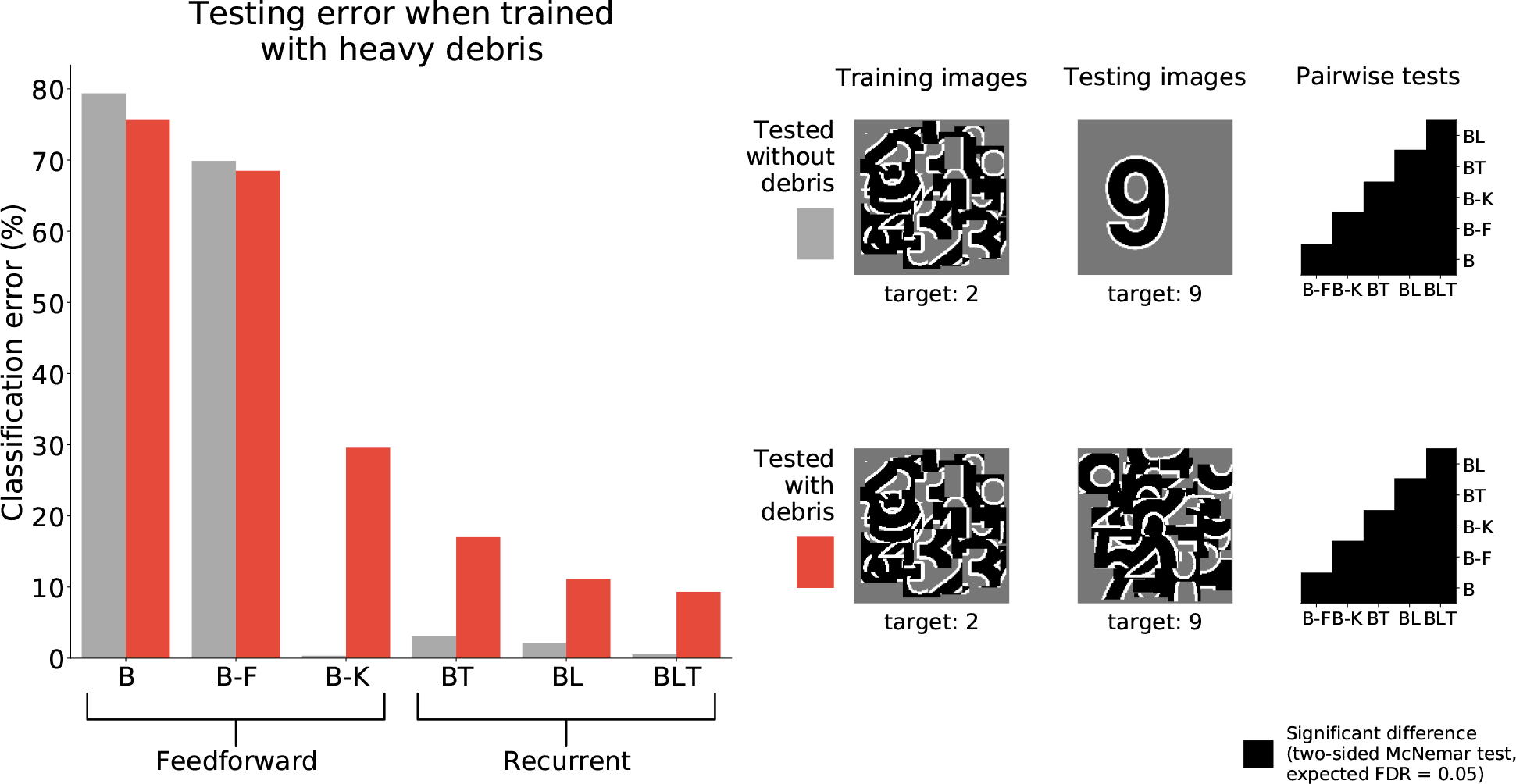
Classification error for all models trained under heavy debris conditions and tested with or without debris. Examples of the images used to train and test the networks are also shown. Matrices to the right indicate significant results of pairwise McNemar tests. Comparisons are across models and within image sets. Black boxes indicate significant differences at p < 0.05 when controlling the expected false discovery rate at 0.05.

**Table 3:** Classification error for all of the models on single digit detection when trained on heavy debris and tested without debris.

| Image set | B | B-F | B-K | BT | BL | BLT |
| --- | --- | --- | --- | --- | --- | --- |
| Tested on debris | 75.63% | 68.49% | 29.58% | 17.01% | 11.13% | 9.32% |
| Tested without debris | 79.37% | 69.88% | 0.34% | 3.10% | 2.11% | 0.55% |

These results show that feedforward networks (specifically B-K) can perform very well at recognising the digit without occlusion, when trained to recognise digits under occlusion. This suggests that they have learnt a good model of the underlying task of digit recognition. However, B-K performs worse than the recurrent models when recognising the target under occlusion. This indicates that B-K has difficulty recognising the digit under occlusion rather than a problem learning to perform the task of digit recognition given the occluded training images. In comparison, recurrent networks show much lower error rates when recognising the target under occlusion.

### 3.2 Recognition of multiple digits

To examine the ability of the networks to handle occlusion when the occluder is not a distractor, the networks were trained and tested on their ability to recognise multiple overlapping digits.

When recognising three digits simultaneously, recurrent networks generally outperformed feed-forward networks (Figure 7), with the exception of BT and B-K where no significant difference was found (*χ*^2^(1, *N* = 30, 000) = 3.53, *p* = 0.06). All other differences were found to be significant (FDR = 0.05). A similar pattern is found when recognising both four and five digits simultaneously. However, in both four and five digit tasks, all pairwise differences were found to be significant, with B-K outperforming BT (Figure 7). This suggests that, whilst recurrent networks generally perform better at this task, they do not exclusively outperform feedforward models.

**Figure 7:**
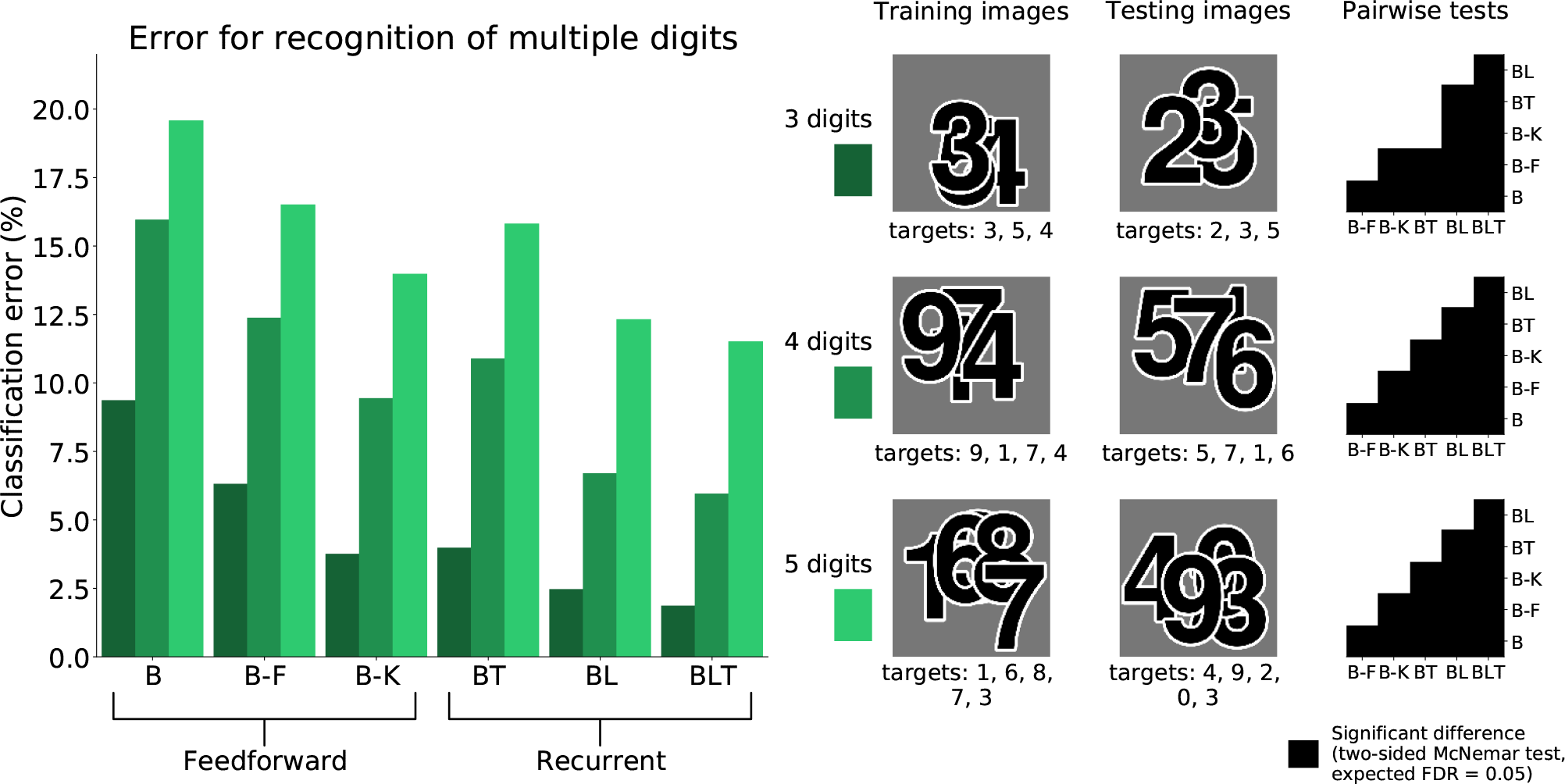
Classification error for all models on multiple digit detection with varying number so of digits. Examples of the images used to train and test the networks are also shown. Matrices to the right indicate significant results of pairwise McNemar tests. Comparisons are across models and within image sets. Black boxes indicate significant differences at p < 0.05 when controlling the expected false discovery rate at 0.05.

### 3.3 MNIST with Gaussian noise

To test the hypothesis that the benefit of recurrence does not extend to dealing with noise in general, we test the performance of the networks on MNIST with unstructured additive Gaussian noise.

The error rates for all models were found to grow as the amount of noise increased (Table 5). Recurrent networks performed significantly better than the feedforward models on MNIST (FDR = 0.05). This supports the idea that recurrent networks are not only better at recognition under challenging conditions, but also in more standard object recognition tasks.

**Table 4:** Classification error for all of the models on multiple digit recognition with varying numbers of targets.

| Image set | B | B-F | B-K | BT | BL | BLT |
| --- | --- | --- | --- | --- | --- | --- |
| 3 digits | 9.35% | 6.30% | 3.74% | 3.97% | 2.45% | 1.85% |
| 4 digits | 15.95% | 12.37% | 9.43% | 10.88% | 6.69% | 5.94% |
| 5 digits | 19.57% | 16.50% | 13.97% | 15.80% | 12.31% | 11.50% |

**Table 5:** Classification error for all of the models on MNIST with varying levels of Gaussian noise.

| Image set | B | B-F | B-K | BT | BL | BLT |
| --- | --- | --- | --- | --- | --- | --- |
| No Noise | 2.99% | 2.42% | 1.43% | 0.95% | 0.95% | 0.95% |
| SNR = 1 | 13.01% | 10.59% | 4.04% | 1.82% | 2.01% | 1.96% |
| SNR = 0.5 | 39.04% | 35.15% | 17.44% | 8.69% | 11.51% | 8.85% |

All pairwise differences were found to be significant between feedforward models. Recurrent networks continued to outperform feedforward networks with the addition of Gaussian noise (Figure 8).

**Figure 8:**
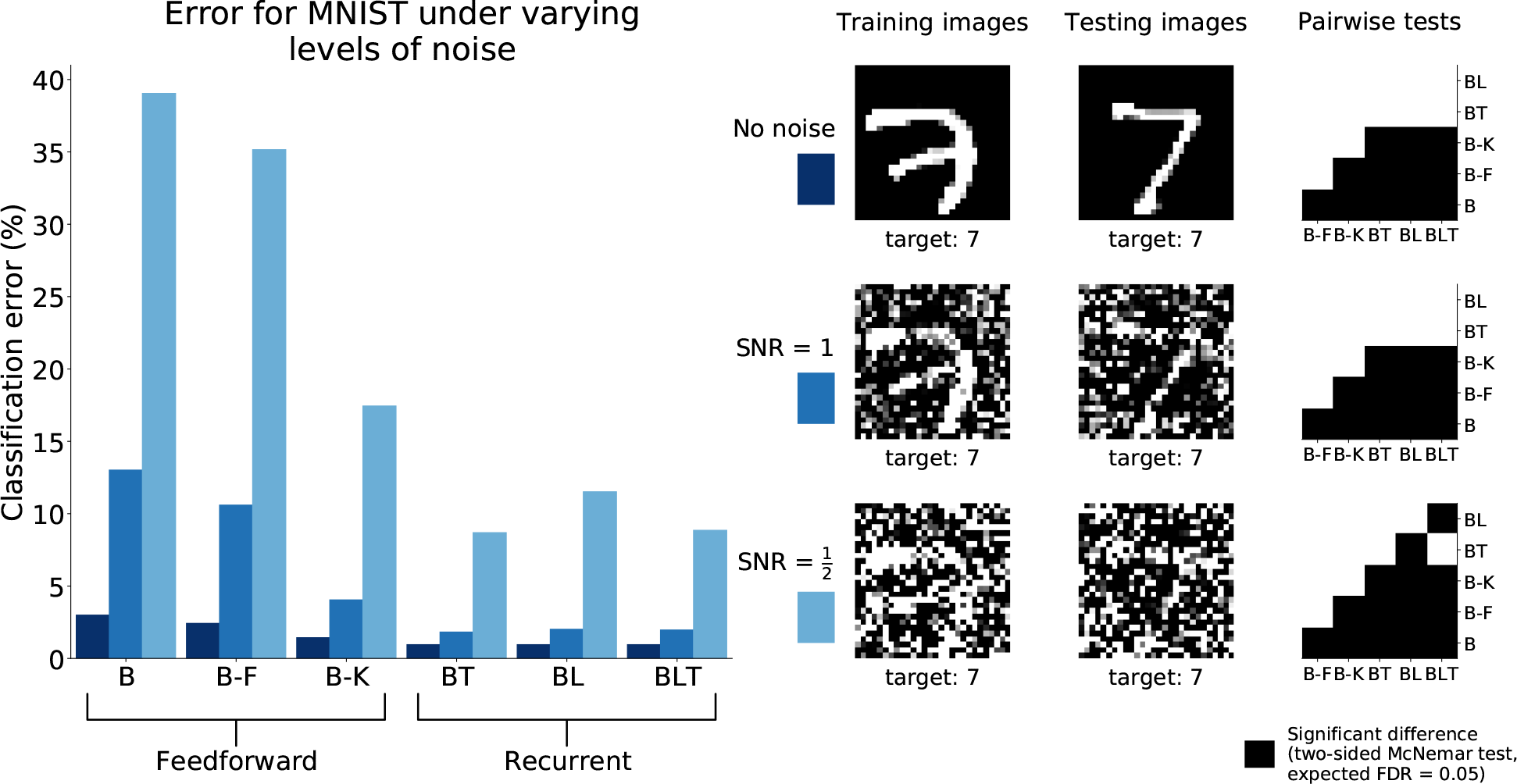
Classification error for all models on recognition in MNIST with and without Gaussian noise. Examples of the images used to train and test the networks are also shown. Matrices to the right indicate significant results of pairwise McNemar tests. Comparisons are across models and within image sets. Black boxes indicate significant differences at p < 0.05 when controlling the expected false discovery rate at 0.05.

At the highest noise levels (SNR = 0.5), BL was found to perform significantly worse than both BT (*χ*^2^(1, *N* = 10, 000) = 61.69, *p* < 0.01) and BLT (*χ*^2^(1, *N* = 10, 000) = 55.12, *p* < 0.01). This means that top-down connections might be more useful for than lateral connections recognising digits under high levels of additive Gaussian noise.

### 3.4 Robustness under challenging conditions

When testing for robustness to increasing levels of debris and Gaussian noise, we found that recurrent networks were always more robust than the feedforward networks. This relationship was not found in the case of clutter. Only one network, BT, was found to be significantly less robust to increases in clutter, and all other networks were found to have similar levels of robustness (Figure 9).

**Figure 9:**
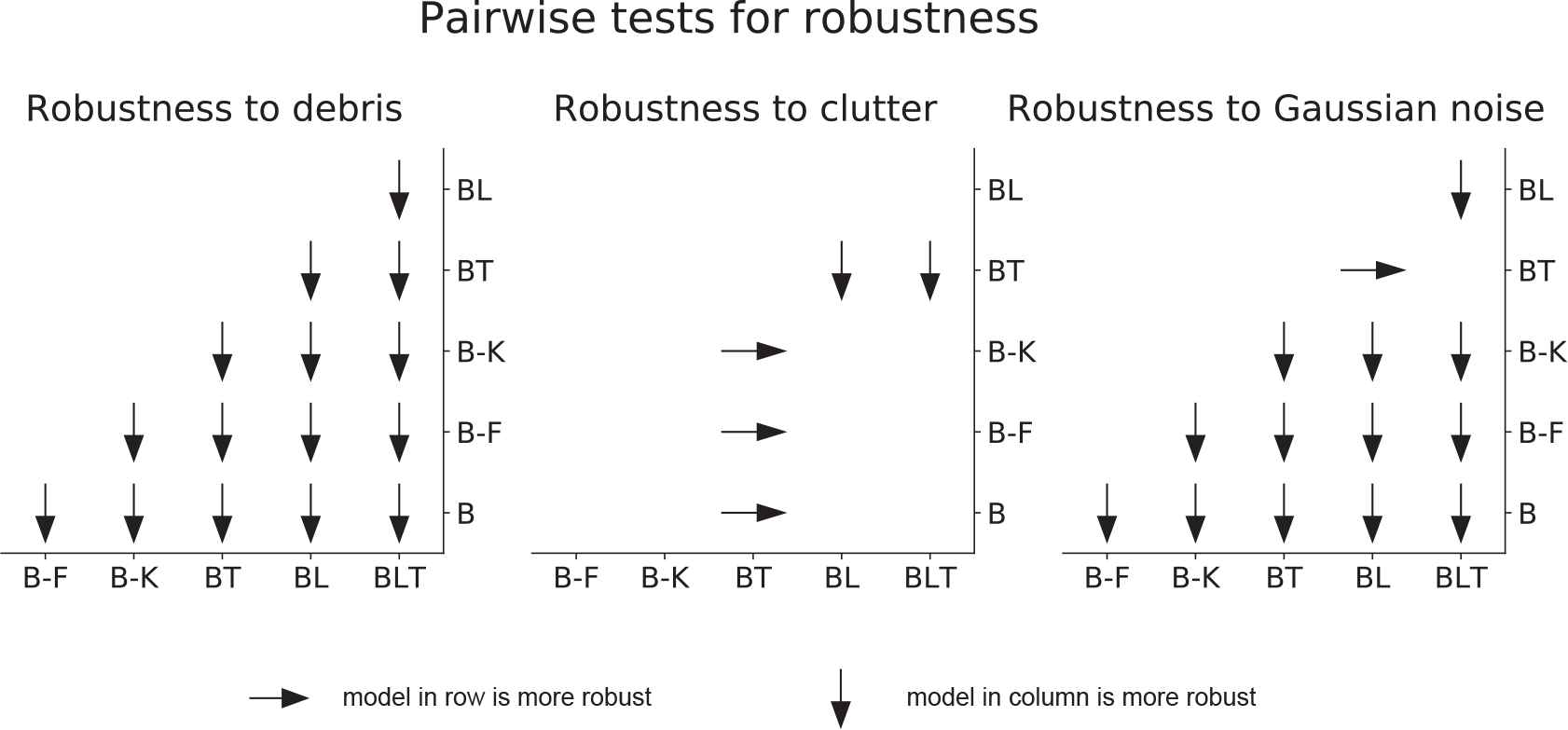
Pairwise differences in model robustness to increased task difficulty. Arrows indicate the more robust model out of the pair tested.

Within feedforward networks, B-K was always the most robust to debris and noise, and B-F was always more robust than B. Within recurrent networks, BLT was the most robust to debris and BL was more robust to debris than BT. However, BLT and BT were more robust than BL to Gaussian noise.

These results suggest that, when debris or Gaussian noise are added, recurrent models take smaller hits to the error rate than feedforward networks. However, when clutter is added, recurrent networks (though still better in absolute performance) take similar hits to the error rate.

More specifically, in the scenarios tested here, lateral recurrence seem to have greater benefit when handling debris and top-down connections improve robustness to Gaussian noise. By utilising both lateral and top-down connections, BLT is more robust to both increasing levels of debris and increasing levels of Gaussian noise.

## 4 Discussion

We found support for the hypothesis that recurrence helps when recognising objects in a range of challenging conditions, as well as aiding recognition in more standard scenarios. The benefit of recurrence for object recognition in challenging conditions appears to be particularly strong in the case of occlusion generated by a non-target and the addition of Gaussian noise, with recurrent networks appearing more robust. In the multiple digit recognition tasks, where the occlusion is generated by other targets, the best performing networks are still recurrent. However, recurrent networks are not more robust, than feedforward networks, to an increased number of digits.

Of the feedforward models, B-K is always the best performing and can outperform recurrent models in some tasks, in the case of multiple digit recognition. One potential explanation is that B-K incorporates some of the benefits of recurrence by having a larger receptive field. This is because recurrence increases the effective receptive field of a unit by receiving input from neighbouring units. This may also explain why BT tends to be the worst performing recurrent model (and outperformed by B-K) in some tasks. BT does not have lateral connections that more directly integrate information from neighbouring units, but information has to go through a higher layer first in order to achieve this. The difference in performance between BT and BL may also tell us about what tasks benefit more directly from incorporating information from outside the classical receptive field (where BL shows an advantage) as opposed to specifically utilising information from more abstract features (where BT shows an advantage). In these experiments, BLT is the best performing network across all tasks, showing that it is able to utilise the benefits of both lateral and top-down connections.

We find evidence to suggest that feedforward networks have particular difficulty recognising objects under occlusion generated by debris, and not just learning the task of recognising digits when trained with heavily occluded objects (Section 3.1.2). This gives specific support to the hypothesis that recurrent processing helps in occluded object recognition.

Recurrent networks also outperformed the parameter matched controls on object recognition tasks where no occlusion was present (Section 3.3). This is consistent with previous work that has shown that recurrent networks, similar in architecture to the BL networks used here, perform strongly compared to other feedforward models with larger numbers of parameters (Liang and Hu, 2015). Therefore, some level of recurrence may be beneficial in standard object recognition, an idea that is supported by neural evidence that shows object information unfolding over time, even without the presence of occlusion (Sugase et al., 1999; Brincat and Connor, 2006; Freiwald and Tsao, 2010; Carlson et al., 2013; Cichy et al., 2014; Clarke et al., 2015).

This work suggests that networks with recurrent connections generally show performance gains relative to feedforward models when performing a broad spectrum of object recognition tasks. However, it does not indicate which of these models best describe human object recognition. Future comparisons to neural or behavioural data will be needed to test the efficacy of these models. For example, as these models are recurrent and unfold over time, they can be used to predict human recognition dynamics for the same stimuli, such as reaction time distributions and the order that digits are reported, in the multiple digit recognition tasks.

Furthermore, we can study whether the activation patterns of these networks predict neural dynamics of object recognition. This is similar to previous work that has attempted to explain neural dynamics of representations using individual layers of deep feedforward networks (Cichy et al., 2016), but by using the recurrent models we can directly relate temporal dynamics in the model to temporal dynamics in the brain. For instance, in tasks with multiple targets (such as those in Section 3.2) we can look at the target representations over recurrent iterations and layers in the model, and compare this to the spatiotemporal dynamics of multiple object representations in neural data. Testing these models against this experimental data will allow us to better understand the importance of lateral and top-down connections, in these models, for explaining neural data.

In addition, whilst we know that adding recurrent connections leads to performance gains in these models, we do not know the exact function of these recurrent connections. For instance, in the case of occlusion, the recurrent connections might complete some of the missing information from occluded regions of the input image, which would be consistent with experimental evidence in cases where parts of the image have been deleted (O'Reilly et al., 2013; Smith and Muckli, 2010). Alternatively, as our occluders contain visual features that could be potentially misleading, recurrent connections may have more of an effect of suppressing the network's representation of the occluders through competitive processing (Adesnik and Scanziani, 2010; Kolankeh et al., 2015). Recurrent connectivity could also learn to produce border ownership cells that would help in identifying occluders in the image (Zhaoping, 2005; Sakai and Nishimura, 2006; Craft et al., 2007), which would help suppress occluders in tasks where occluders are non-targets. If these networks are to be useful models of visual processing, then it is important that future work attempts to understand the underlying processes taking place.

It could be argued that BLT performs the best due to the larger number of parameters it can learn. However, we know that the performance of these networks is not only explained by the number of learnable parameters, as B-F has the largest number of parameters of the models tested (Table 1) and performs poorly in all tasks relative to the recurrent models. Finding exactly parameter matched controls for these models that are conceptually sound is difficult. As discussed earlier (Section 2.2), altering the kernel size of the feedforward models is the best control, but this provides a relatively coarse-grained way to match the number of parameters. Altering the number of learnt features allows more fine-tuned controls for the number of parameters, but this also changes the number of units in the network, which is undesirable. We believe that the models used here represent a good compromise between exact parameter matching and the number of units in each model.

This research suggests that recurrent convolutional neural networks can outperform their feed-forward counterparts across of diverse set of object recognition tasks and that they show greater robustness in a range of challenging scenarios, including occlusion. This builds on previous work showing a benefit of recurrent connections in non-convolutional networks where parts of target objects are deleted (O'Reilly et al., 2013). This work represents initial steps for using recurrent convolutional neural networks as models of visual object recognition. Scaling up these networks and training them on large sets of natural images (e.g. Russakovsky et al., 2015) will also be important for developing models that mirror processing in the visual system more closely. Future work with these networks will allow us to capture temporal aspects of visual object recognition that are currently neglected in most models, whilst incorporating the important spatial aspects that have been established by prior work (DiCarlo et al., 2012). Modelling these temporal properties will lead to a more complete understanding of visual object recognition in the brain.

## Acknowledgements

This research was funded by the UK Medical Research Council (Programme MC-A060-5PR20), by a European Research Council Starting Grant (ERC-2010-StG 261352), and by the Human Brain Project (EU grant 604102 ‘Context-sensitive multisensory object recognition: a deep network model constrained by multi-level, multi-species data’).

